# Methylation of Q105 on histone H2A is part of a dynamic regulatory mechanism integrating metabolism with ribosome biogenesis through recruitment of Nhp2

**DOI:** 10.1101/2021.01.11.426220

**Authors:** Julia S.P. Mawer, Niklas Grabenhorst, Constantine Mylonas, Peter Tessarz

## Abstract

Ribosome biogenesis is an essential cellular process that requires integration of extracellular cues, such as metabolic state, with intracellular signaling, transcriptional regulation and chromatin accessibility at the ribosomal DNA. Here, we demonstrate that the recently identified histone modification, methylation of H2AQ105, is an integral part of a dynamic chromatin network at the rDNA locus. Its deposition depends on a functional mTor signaling pathway as well as acetylation of histone H3 at position K56, thus integrating signals from cell cycle, metabolic and proliferative states. Furthermore, we identify a first epigenetic reader of this modification, the ribonucleoprotein Nhp2, which specifically recognizes the methylation on H2AQ105. Based on functional and proteomic data we suggest that Nhp2 functions as an adapter to bridge the rDNA chromatin with components of the small subunit processome and might help to efficiently coordinate transcription of rRNA with its post-transcriptional processing.

## INTRODUCTION

Regulation of ribosomal biogenesis is a complex process. It requires all three eukaryotic polymerases and is tightly linked to the cell cycle (Bernstein *et al*, 2007; Piazzi *et al*, 2019) and metabolic state, particularly via Tor-mediated signaling (Mayer & Grummt, 2006; Huber *et al*, 2011). It was noted early on that inhibition of Tor kinase by rapamycin leads to shut-down of ribosomal RNA (rRNA) transcription by RNA polymerase I (RNA PolI) and concomitant decrease in the levels of acetylation of lysine 5 and 12 on histone H4, demonstrating that chromatin plays an important role in the regulation of RNA PolI, but also can also be used to read-out its activity based on the chromatin profile (Tsang *et al*, 2003). H3K56 acetylation is another modification that has been described to be dependent on Tor signaling (Chen *et al*, 2012). In particular, histone acetylation seems to be sensitive towards inhibition by rapamycin - potentially as it marks actively transcribed chromatin.

Transcription of rRNA is followed by several post-transcriptional processing steps including base and ribose methylation or pseudouridylation of the rRNA. Base methylation occurs late in ribosome maturation and only in highly conserved rRNA sequences. Pseudouridylation is thought to play a variety of roles in the ribosome, including the improvement of translational fidelity (Jack *et al*, 2011). The majority of ribose methylation takes place early in rRNA processing and is considered to be important for rRNA folding or association with chaperone proteins that might help folding of rRNA. Single sites of ribose methylation are not essential (Weinstein & Steitz, 1999), but global rRNA demethylation severely impairs growth (Tollervey *et al*, 1993).

Pioneering studies using electron microscopy demonstrated that rRNA folds co-transcriptionally (Scull & Schneider, 2019). Importantly, RNA Pol I transcription speed and folding kinetics are directly coupled as shown by mutations in RNA Pol I that decrease the rate of synthesis (Schneider *et al*, 2007). Not only does lower transcription speed impact rRNA folding, but it also directly affects rRNA processing (Duss *et al*, 2018). In addition, previous work showed that early processing steps, particularly of the small ribosomal subunit components also occur co-transcriptionally (Gallagher *et al*, 2004; Kos & Tollervey, 2010).

Taken together, these data suggest a tight link between chromatin architecture, transcriptional regulation, synthesis rate and processing. One potential way of linking these different steps in an efficient way would be to directly recruit the processing machinery to the rDNA chromatin. We previously identified methylation of H2AQ105 as an RNA Pol I dedicated histone modification that is exclusively enriched at the rDNA locus (Tessarz *et al*, 2014). The modification is dependent on RNA PolI-mediated transcription and is catalyzed by the nucleolar methyltransferase NopI/Fibrillarin (Tessarz *et al*, 2014; Loza-Muller *et al*, 2015; Iyer-Bierhoff *et al*, 2018).

Here we show that H2AQ105me is an integral part of a dynamic chromatin network at the rDNA locus. Deposition of the modification in yeast is cell cycle dependent - similar to human cells (Iyer-Bierhoff *et al*, 2018) - and relies on proliferation. Furthermore, we can demonstrate that the methylation is downstream of the Tor signaling pathway and acetylation of histone H3 at position K56. We identify Nhp2 as an epigenetic reader of H2AQ105me and provide evidence that Nhp2 might function as an adapter to bridge chromatin and components of the SSU.

## RESULTS

### H2AQ105 methylation is a dynamic histone modification that requires cell proliferation

As ribosome biogenesis is tightly linked to cellular state, we initially sought to determine whether H2AQ105me levels, in the yeast *Saccharomyces cerevisiae*, were regulated in a cell-cycle dependent manner. Cells were arrested in G1 using alpha factor. Following release of G1 arrest, cells were sampled at time intervals and levels of H2AQ105me were assessed by western blotting (Figure 1A). H2AQ105me levels fluctuate throughout the cell cycle and peak in S-phase, as shown by the co-increase of H2AQ105me and the S-phase marker, H3K56ac. This data confirms observations in human cells, in which a cell cycle-dependent fluctuation of H2AQ105me was reported (Iyer-Bierhoff *et al*, 2018). Interestingly, arresting the cell cycle in any phase leads to loss of H2AQ105me, indicating that proliferation is required for the methylation (Figure 1B). Even an arrest in S-phase using the drug hydroxyurea (HU), which depletes the cell of deoxynucleotides leading to DNA replication fork stalling and collapse, leads to loss of Qme rather than its accumulation, as is seen for H3K56ac (Figure 1B). This observation is likely explained by a recent study in which rDNA transcription dynamics were studied under HU-induced cell-cycle arrest, showing that rDNA transcription, as well as DNA replication, is halted by HU (Charton *et al*, 2019).

**Figure 1:**
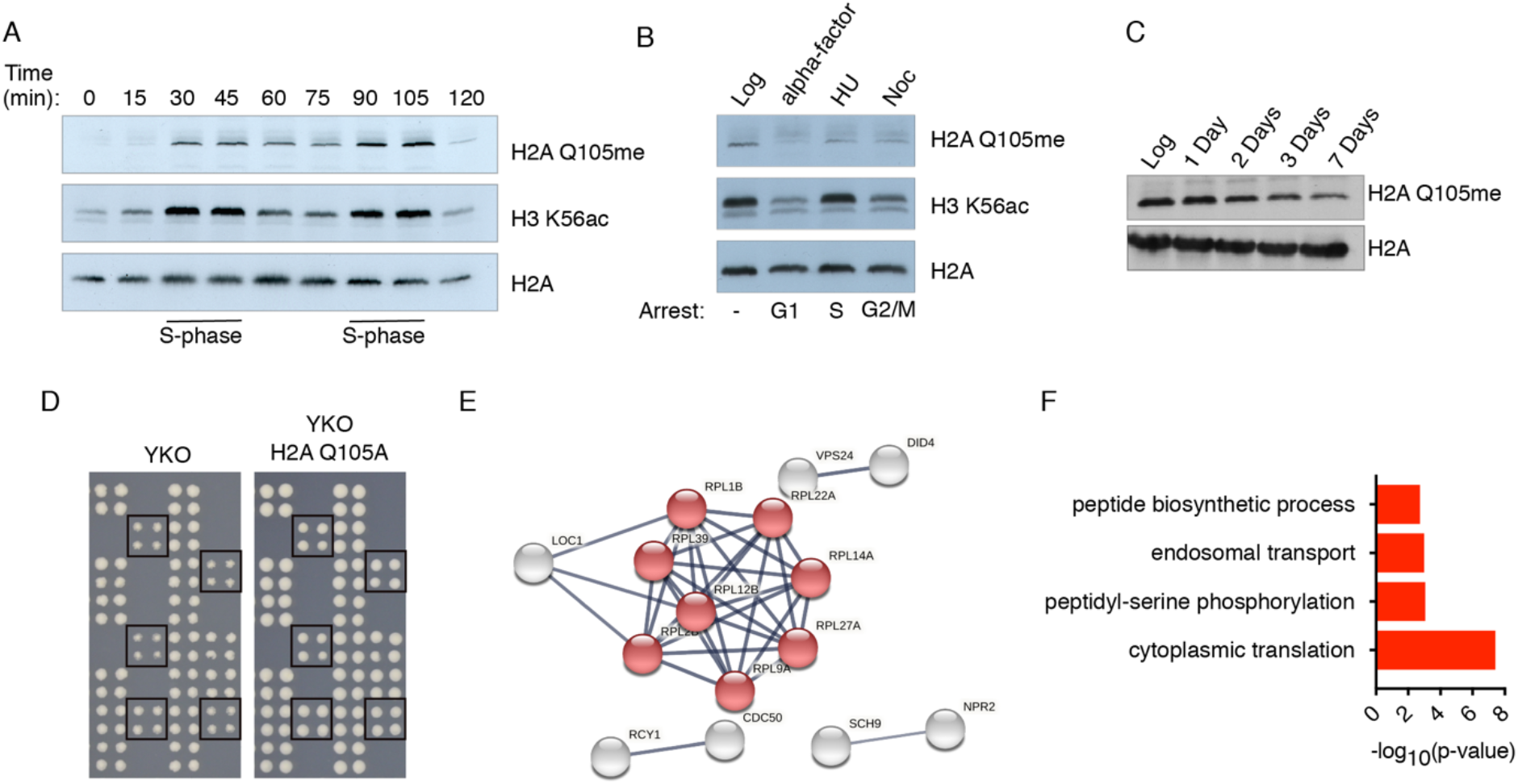
H2AQ105me fluctuates with the cell cycle and is a mark of proliferating cells. A) Cell cycle profile of H2AQ105me. Cells were arrested using alpha-factor. Upon release (t=0), samples were taken every 15 minutes. S-phase is indicated by the enrichment of H3K56ac. B) H2AQ105me levels upon arrest at different cell cycle stages. Treatment is giving above the lanes. C) H2AQ105me levels upon growth of cells into stationary phase. D) Example of an SGA plate highlighting rescue of slow growth, E) STRING network of genes identified in SGA and F) GO enrichment for these genes.

This strengthens the observation that RNA PolI-mediated transcription of the rDNA is required for deposition of H2AQ105me and that the methylation is associated with transcribed rDNA repeats. To explore the requirement for proliferation further, yeast cells were grown deep into stationary phase (Figure 1C) by growing in a sealed flask for up to one week without replenishing the growth medium or diluting the density of cells. Cells were sampled during log-phase of growth at days 1, 2, 3, and 7. Western blot analysis shows a progressive decrease in levels of H2AQ105me over time.

In order to identify pathways that might be responsible for the regulation of H2A105 methylation, we performed a synthetic genetic array (SGA) (Tong *et al*, 2001), by incorporating an H2AQ105A mutant into the genome, which was subsequently mated against the full knock-out collection. We did not observe any negative genetic interaction for H2AQ105A, but noted that the growth of several slow growing deletion strains was rescued upon combination with H2AQ105A (Figure 1D and Supplementary Table 1). Intriguingly, many ribosomal genes were identified, as were members of the mTor signaling pathway, such as Sch9, the yeast homolog of S6 kinase (Figure 1E, F).

### Methylation of H2AQ105 is regulated by the mTor signaling pathway and acetylation of H3K56

The observation that an alanine substitution at H2AQ105 could rescue the slow growth of an *sch9* deletion in the SGA led us to investigate the role of mTor in the methylation of H2AQ105. It is well known that mTor is a critical regulator of rDNA transcription (Mayer & Grummt, 2006). Indeed, addition of rapamycin led to a loss of RNA PolI from the rDNA locus (Supplementary Figure 1A) and a stop in proliferation (Supplementary Figure 1B). Interestingly, mTor has also been demonstrated to be also involved in the regulation of H3K56 acetylation at the rDNA locus (Chen *et al*, 2012), a histone modification that cycles with H2AQ105me as part of the cell cycle (Figure 1A). We confirmed the described decrease in H3K56 acetylation upon inhibition of mTor using Rapamycin at the rDNA by chromatin immunoprecipitation (ChIP) (Chen *et al*, 2012) (Figure 2B,C). We then performed ChIPs using an H2AQ105me specific antibody (Tessarz *et al*, 2014). As with H3K56ac, we observed a time-dependent loss of H2AQ105me, which was restricted to the coding region of the rDNA (Figure 2D, E). Interestingly, in both cases, we did not observe an increase in the deposition of the core histone (Figure 2B, D). Given the similarities in the kinetics of reduction for both investigated histone marks, we wondered if there was a connection between these two modifications. Mutating H3K56 to alanine led to a strong reduction of H2AQ105me (Figure 2F). Importantly, mutations at other histone side chains, also modified in a cell cycle-dependent manner, did not have an effect on the level of H2AQ105me (Figure 2F). To corroborate the dependence of H2AQ105me on H3K56ac, we went on and analyzed the machinery involved in the acetylation of H3K56. Before deposition into chromatin, H3 is bound by the histone chaperone Asf1 (Chen *et al*, 2008). The histone acetyl-transferase Rtt109 subsequently binds the Asf1-H3/4 complex and acetylates H3 at K56 (Chen *et al*, 2008) prior to the incorporation of the H3/4 tetramer into chromatin. Deletions of both *asf1* and *rtt109*, led to a reduction of H2AQ105me confirming the result of the mutational analysis in H3 (Figure 2G). Rtt109 is a slow growing strain and this reduction in growth might be responsible for the reduction in H2AQ105me. However, growth is rescued by the additional mutation in H3K56A (Han *et al*, 2007), while H2AQ105me levels remain low (Figure 2G), indicating that acetylation at H3K56 is an important contributor for H2AQ105 methylation. H3K56ac is de-acetylated by Hst3/4 (Kaplan *et al*, 2008). In line with the idea that acetylation of H3K56 is required for H2AQ105me, deletions of *hst3/4* did not change the level of H2AQ105me (Figure 2G). Finally, we wanted to address how acetylation of H3K56 mediates H2AQ105me. An alanine substitution at H3K56 as well as an *asf1* deletion led to a reduction of PolI occupancy at the rDNA (Chen *et al*, 2012). Thus, it is plausible that the observed dependency of H2AQ105me on H3K56ac is due to a reduction in transcription rate and a concomitant decrease in Nop1 recruitment to the rDNA. To test this, we used a FLAG-tagged version of Nop1 and performed ChIP using a FLAG antibody. Indeed, Nop1 was efficiently enriched over the transcribed region of the rDNA locus, but this enrichment was decreased in the presence of an alanine substitution at H3K56 (Figure 2H). Taken together, methylation of H2AQ105me depends on active rDNA transcription, H3K56 acetylation and the integration of metabolic cues via the mTor pathway.

**Figure 2:**
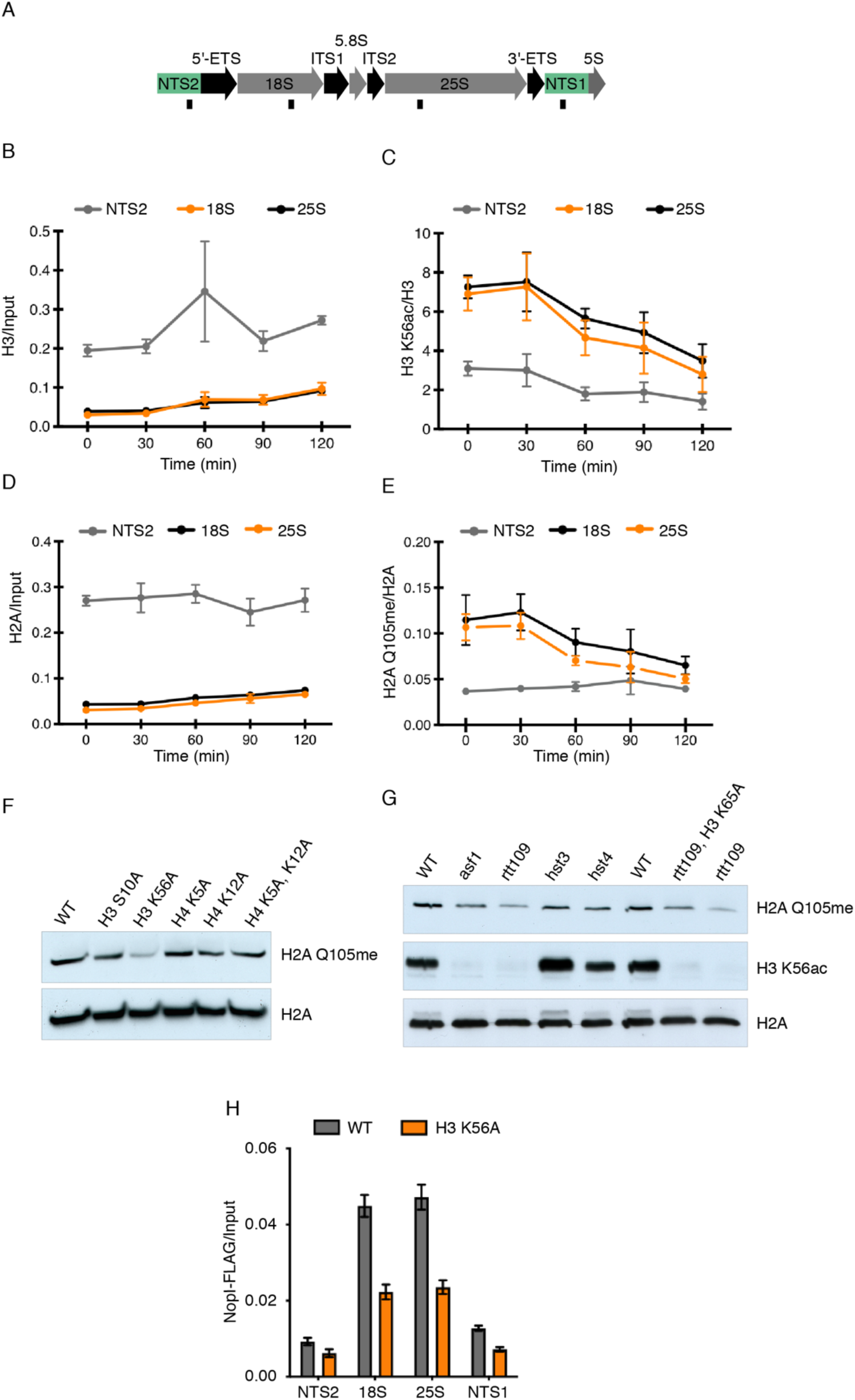
H2AQ105me is regulated by Tor signaling and H3K56 acetylation. A) Scheme of rDNA locus and indication of primer locations used in B-E. B-E) ChIP-qPCR for indicated antibodies after treatment with 5nM rapamycin. Samples were either normalized to input or to the core histone. F) H2AQ105me levels in various histone mutations targeting sites of well-known cell cycle-dependent modifications. G) H2AQ105me and H3K56ac levels in deletions of genes important for H3K56ac deposition. H) ChIP-qPCR for Nop1-FLAG using anti-FLAG tag antibody. Samples were normalized using input.

### Ribosome biogenesis factors are enriched on H2AQ105me

Previous work identified the region surrounding H2AQ105 as part of a recognition sequence for the histone chaperone FACT (McCullough *et al*, 2011). Methylation of Q105 prevents FACT from binding to H2A, impairing its redeposition following transcription and resulting in loss of histones at the rDNA (Tessarz *et al*, 2014). Given the link of H2AQ105me with active rRNA transcription and its demonstrated role to inhibit binding, we next wanted to address if methylation of H2AQ105 could also serve as a recognition site for the recruitment of protein complexes to the rDNA. As this histone modification is exclusively localised in the nucleolus (Tessarz *et al*, 2014; Loza-Muller *et al*, 2015; Iyer-Bierhoff *et al*, 2018), it would be ideally suited to serve as a chromatin beacon to signal actively transcribing RNA PolI.

We used a peptide pulldown approach that has been successfully performed on several histone modifications (Vermeulen *et al*, 2010). Following SILAC labelling of yeast cultures, whole cell lysis pulldowns were performed using unmodified and modified H2A peptides spanning the region of Q105 and analysis was performed by quantitative mass spectrometry (Figure 3A). Importantly, we observed Pob3 and Spt16 - both subunits of the yeast FACT complex - to be enriched on the unmodified H2A peptide (Figure 3B). 139 proteins were found to be significantly, 2-fold or more, enriched on peptides harbouring the modification (Figure 3B and Supplementary Table 2). Analysis of this list using the online protein-protein interaction network and functional analysis tool, String (https://string-db.org/), showed that these proteins form a tight network (Figure 3B). Gene ontology (GO) enrichment highlighted this network to be composed of ribosome biogenesis factors and ribosomal proteins (Figure 3C). These data suggest that H2AQ105me might indeed help to recruit ribosome biogenesis factors to the site of active rDNA transcription. As much of the rRNA processing occurs co-transcriptionally (Turowski & Tollervey, 2015), recruitment of processing factors to the rDNA chromatin would facilitate this process.

**Figure 3:**
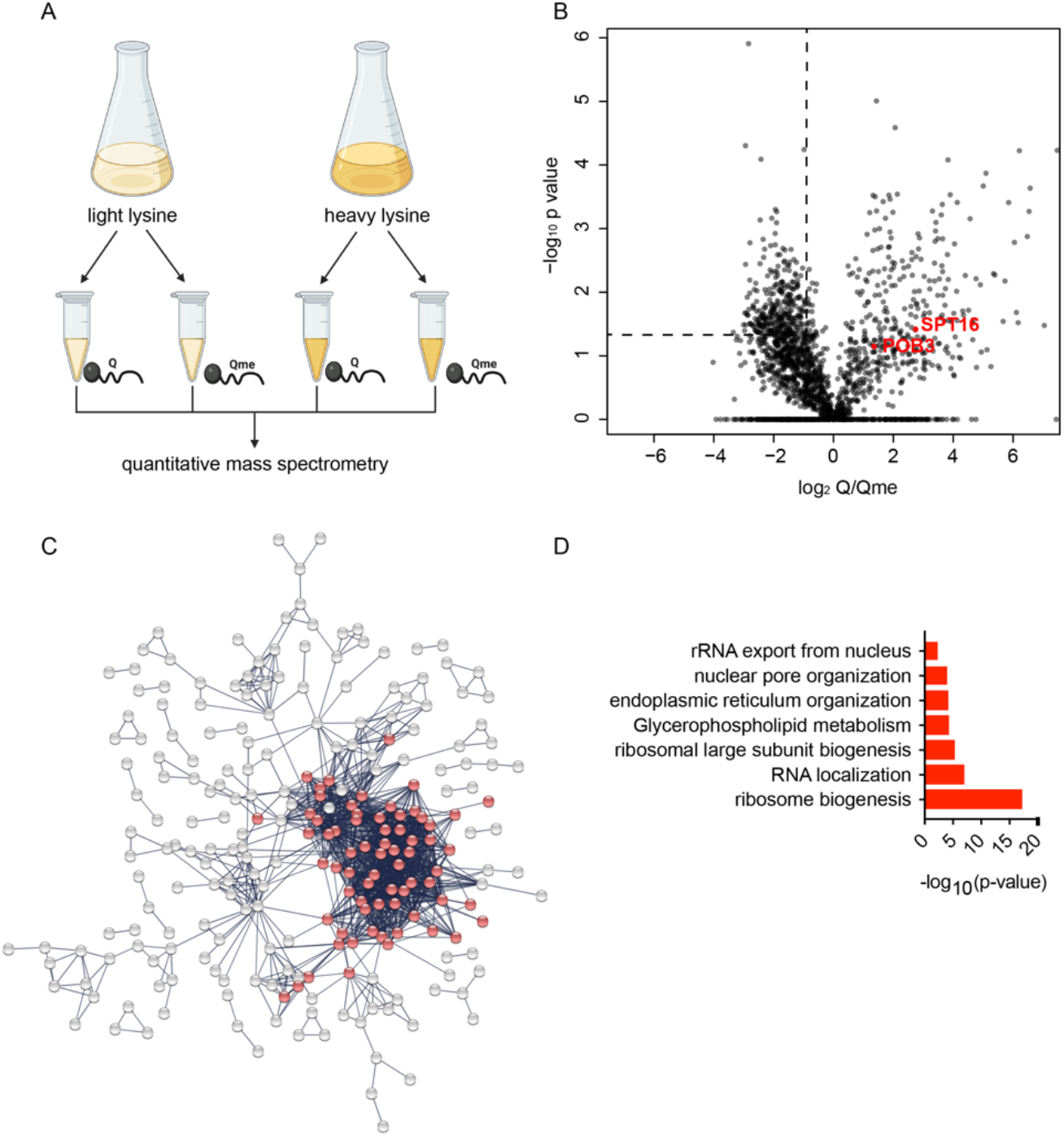
Mass spectrometry identifies ribosome biogenesis factors to be recruited to H2AQ105me. A) Schematic of the experimental strategy. B) Volcano plot of quantitative SILAC experiments. Left upper quadrant indicates significantly, 2-fold enriched proteins bound specifically to H2AQ105me. Members of the FACT complex (SPT16/POB3) are highlighted in red. C) STRING network of proteins enriched on H2AQ105me (highlighted in red are proteins involved in ribosome biogenesis). Protein names can be found in Supplementary Table 2. D) GO enrichment for proteins bound to H2AQ105me.

### Nhp2 is a reader of H2AQ105me

In order to gain further clarity on a possible mechanism for this recruitment, it was important to identify a direct binder of glutamine methylation, an epigenetic reader of this histone modification. In order to do this, candidates were selected from the list of 139 Qme-enriched proteins identified by mass spec (Figure 3B), based on their known interactions listed on the yeast genome database (https://www.yeastgenome.org/). Initially five potential readers were myc-tagged to allow for their easy detection and ChIPs were performed in wildtype and H2AQ105A strains. Only one of these five candidates - Nhp2 - was efficiently enriched on the rDNA, while recruitment was decreased in an H2AQ105A background (Figure 4A and Supplementary Figure 2). We then repeated the peptide pulldowns from yeast carrying the myc-tagged Nhp2 allele. Nhp2 showed enrichment on the peptide harbouring the methylation on Q105, confirming the mass spectrometry data (Figure 3B) and highlighting Nhp2 as a possible epigenetic reader of H2AQ105me. To confirm this hypothesis, Nhp2 was recombinantly purified from *Escherichia coli*. Peptide pulldowns using only recombinant Nhp2 were performed and analysed by SDS-PAGE and Coomassie staining (Figure 4C). This approach was able to confirm the ability of Nhp2 to directly bind to the peptide containing the methylated glutamine (Figure 4C).

**Figure 4:**
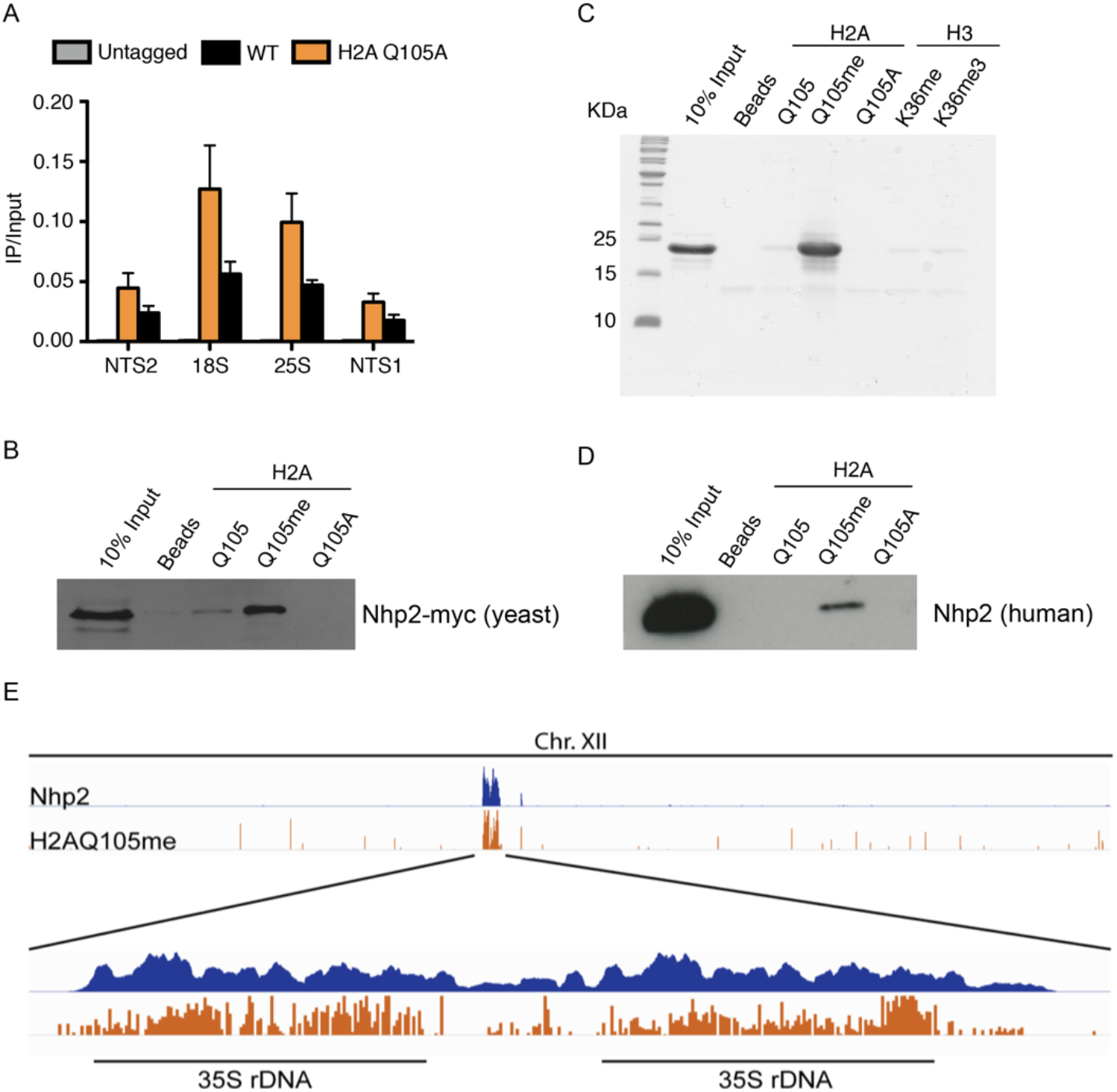
Nhp2 is an epigenetic reader of H2AQ105me. A) ChIP-qPCR for Nhp2-myc using anti-myc antibody across the rDNA locus. Samples were normalized to input. B) Peptide pulldown on H2AQ105 peptides from yeast expressing myc-tagged Nhp2. C) Peptide pulldown using recombinantly purified Nhp2. D) Peptide pulldown on H2AQ105 peptides from human cells (HEK293T). E) ChIP-seq profile of Nhp2 (blue) and H2AQ105me (orange, data from Tessarz *et al*., 2014) across chromosome XII of yeast. Magnification shows the two annotated rDNA loci in the yeast genome.

Furthermore, Nhp2 did not bind to methyl-containing peptide sequences derived from histone H3, further confirming its specificity to methylation of Q105 on histone H2A (Figure 4C). H2AQ105me is conserved from yeast to mammalian cells (Tessarz *et al*, 2014) - as is Nhp2. To test if the Nhp2-H2AQ105me interaction might also be evolutionarily conserved, we performed peptide pulldowns using mammalian cell lysates. While the interaction was not as strong as in the case for yeast, human Nhp2 clearly interacts specifically with methylated H2AQ105 (Figure 4D), confirming the evolutionary conservation of the Nhp2-H2AQ105me interaction. Finally, we wanted to address whether Nhp2 recruitment would follow H2AQ105 methylation genome wide. We performed ChIP-sequencing using Nhp2-myc. In line with the data described above, the ChIP profile of Nhp2 closely resembles the one of H2AQ105me with a strong enrichment over the rDNA region on yeast chromosome XII (Figure 4E), corroborating the idea that Nhp2 is an epigenetic reader of methylated H2AQ105.

### Nhp2 bridges H2AQ105me to ribosome biogenesis factors

Nhp2 is a highly conserved protein that has been identified to be a member of the H/ACA ribonucleoprotein complex and binds snoRNAs. Together with Gar1, Nop10 and the catalytic subunit, Cbf5, it is responsible for the pseudouridylation of ribosomal RNA (Li *et al*, 2011b; Wang & Meier, 2004; Fath *et al*, 2000). Nhp2 is an essential gene and is required for the maintenance of snoRNA levels (Henras *et al*, 1998). On the contrary, pseudouridylation of rRNA is not essential for yeast, but is required to maintain translational fidelity (Jack *et al*, 2011). Interestingly, pseudouridylation has been described to potentially occur co-transcriptionally (Penzo & Montanaro, 2018). Yeast strains lacking pseudouridylation show an increase in frameshifts and are affected in tRNA binding to the ribosome (Jack *et al*, 2011). This can be easily measured by the sensitivity of yeast strains to several antibiotics that bind the A- and P-sites of the ribosome. Strains lacking pseudouridylation are hyper-sensitive to paromomycin and show higher resistance against anisomycin (Jack *et al*, 2011). While the hypothesis of H2AQ105me-dependent recruitment of the pseudouridylation machinery was an attractive hypothesis, it is important to note that we did not detect an enrichment for any of the other components of the pseudouridylation complex using peptide pulldowns (Supplementary Table 2 and Figure 3B). Additional pulldowns using TAP-tagged components of this complex confirmed the mass spectrometry data (Figure 5A). Indeed, we did not observe any change in sensitivity towards paromomycin or anisomycin in H2AQ105A strains (Figure 5B), indicating that pseudouridylation is not strongly affected in the absence of H2AQ105-methylation. To confirm this, we used ribosome translational slippage reporters as a more sensitive assay for changes in pseudouridylation status of the rRNA (Jack *et al*, 2011). In these reporters, the firefly and renilla luciferases are separated by sequences known to cause translational frameshift mutations.

**Figure 5:**
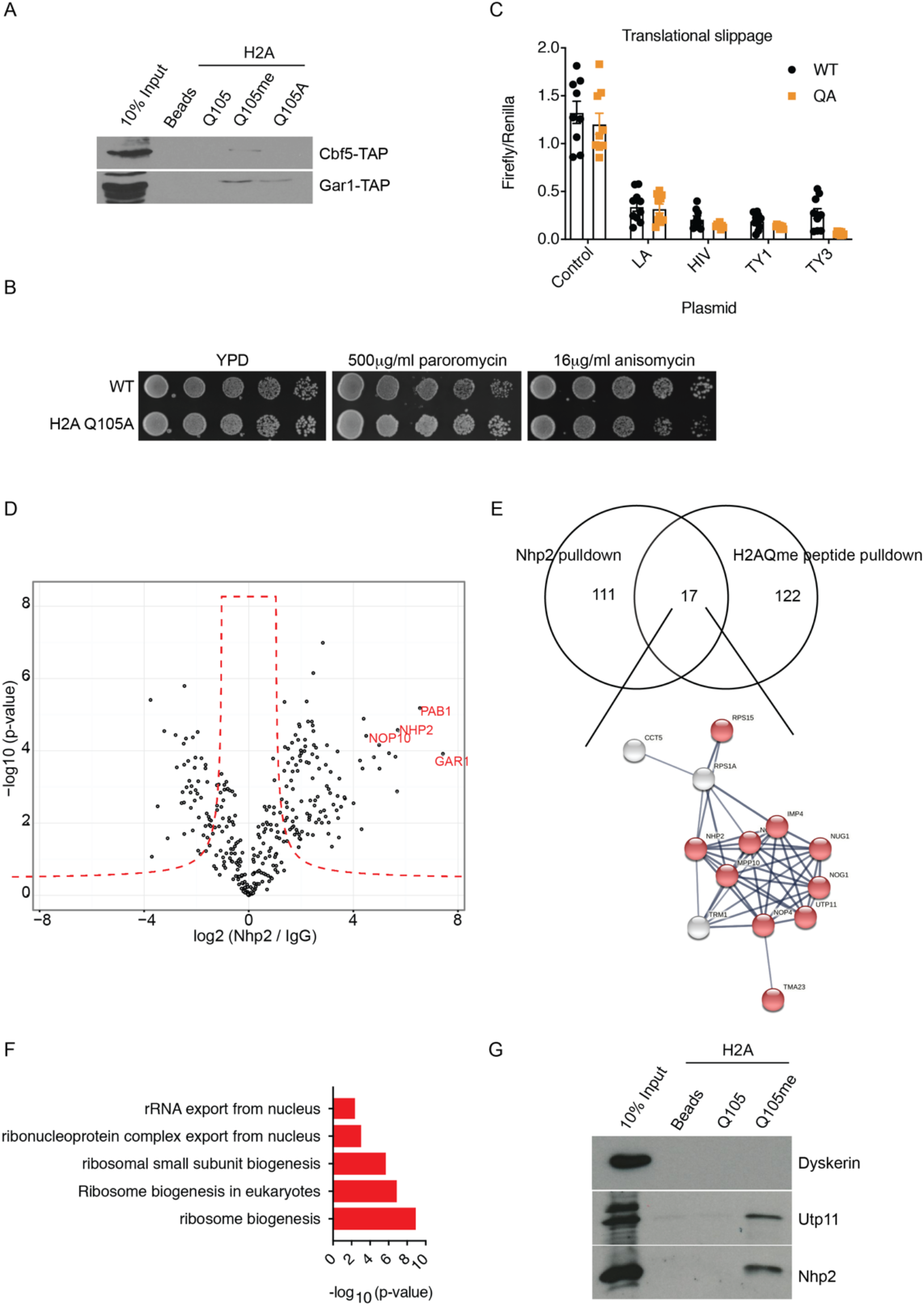
Nhp2 bridges components of the SSU and H2AQ105me. A) Peptide pulldowns from indicated TAP-tagged strains. Pulldown efficiency was analyzed by western blotting. B) Spot test of the indicated strains on YPD or YPD supplemented with the given antibiotics. C) Ribosome slippage assay using a tandem luciferase reporter. The sequences allowing for slippage are given below the graph. D) Proteomic analysis of Nhp2 interactors. Highlighted in red are the known members of the H/ACA pseudouridylation complex. E) Intersection of Nhp2 and H2AQ105me pulldown. 17 proteins were enriched in both experiments and the majority are members of the small subunit processome (highlighted in red). F) GO enrichment for these 17 commonly identified proteins. G) Peptide pulldowns from human cells (HEK293T) showing conservation of the reported interactions in yeast. Pulldown efficiency was analyzed by western blotting using the indicated antibodies.

When this frameshift occurs, the translation of the second luciferase is induced (Harger & Dinman, 2003). No significant difference in translational frameshift was observed between a wildtype and H2AQ105A strains (Figure 5C). Given that we were not able to identify evidence for a link between H2AQ105me and pseudouridylation, we considered that Nhp2 might have additional functions outside the H/ACA pseudouridylation complex. In order to determine the role that Nhp2 binding to H2AQ105me could play, we performed immunoprecipitation (IP) followed by quantitative mass spectrometry using the myc-tagged version of Nhp2. Enrichment was calculated against the untagged parental strain. 128 proteins were found to be significantly, 2-fold or more, enriched by myc in the Nhp2-myc strain (Figure 5D). Among the top enriched proteins were all members of the pseudouridylation complex, indicating that the IP worked successfully (Figure 5D). To identify proteins that might be recruited to H2AQ105me via Nhp2, we intersected the list of Nhp2 binders with the proteins recruited to H2AQ105me (Figure 3B). 17 proteins were enriched in both experiments (Figure 5E). String analysis identified again a tight network of interacting proteins (Figure 5E). Gene ontology enrichment revealed these to be ribosome biogenesis factors, particularly involved in the processing of the small ribosomal subunit, like Mpp10, Imp4 or Utp11 (Figure 5F). Finally, we tested if this might be conserved in humans as human Nhp2 interacted with H2AQ105me (Figure 4D). Therefore, we tested if Dyskerin (the human Cbf5 homolog) or Utp11 (member of the SSU) would interact with H2AQ105me. In line with the observations from yeast, Dyskerin was not pulled down on H2AQ105me-containing peptides, while Utp11 was enriched to a similar extend as Nhp2 (Figure 5G), indicating that Nhp2 might indeed serve as an adapter protein involved in the recruitment of factors involved in the early steps of ribosome maturation.

## DISCUSSION

A chromatin mark that signals actively transcribing rRNA might be an ideal way for cells to integrate proliferation and signaling with ribosome biogenesis, while at the same time maintain chromatin architecture for an efficient transcription. Recently, it was shown that phosphorylation of tyrosine 11 in histone H3 was an endpoint of a signaling cascade that integrates a response to nutrient availability and ultimately, shaped chromatin to allow efficient transcription (Oh *et al*, 2018). One signaling branch leading to H3T11ph leads via Tor and Sch9, the yeast homolog of S6 kinase. However, H3T11ph is found genome-wide, including metabolic and ribosomal protein genes, as well as rDNA (Oh *et al*, 2018). The recently discovered histone modification H2AQ105me (Tessarz *et al*, 2014) might represent a mark that works exclusively on rDNA and thus, could integrate metabolic and proliferation cues directly with rDNA transcription. Here, we show that H2AQ105me is regulated via Tor signaling and inhibition of Tor by rapamycin leads to loss of methylation across the rDNA. The methylation also depends on the proper incorporation of H3K56 acetylated histone H3 at the rDNA (Figure 2). However, it is not clear if the links between Tor signaling and H3K56 acetylation with H2AQ105me are direct or a consequence of decreased transcription at the rDNA. In mammalian cells, methylation of H2AQ105me directly depends on the acetylation status of Fibrillarin, which regulates binding to H2A and appears to be independent of the transcriptional output of the rDNA (Iyer-Bierhoff *et al*, 2018). The cycling of H2AQ105me with the cell cycle (Figure 1 and (Iyer-Bierhoff *et al*, 2018)) and the dependence of the methylation on H3K56 acetylation point towards a role for H2AQ105me in the re-establishment of efficient transcription at the rDNA upon DNA replication and thus might be a great cellular reporter for proliferative cells.

A chromatin modification that specifically marks the rDNA might also be a recruitment site for factors involved in the co-transcriptional processing of rRNA. Many of the early processing steps, particularly methylations and processing of the 18S rRNA have been shown to occur co-transcriptionally (Gallagher *et al*, 2004; Kos & Tollervey, 2010). Here, we identified Nhp2 to be a direct reader of H2AQ105 methylation. Nhp2 is an RNA binding protein that binds a multitude of small nuclear RNAs. Depletion of Nhp2 leads to a strong decrease in the levels of many snoRNAs and this appears to be the reason why Nhp2 is essential (Henras *et al*, 2001). Nhp2 recruits snoRNAs to the H/ACA pseudouridylation complex that are subsequently used as guides to position the catalytic subunit, Cbf5, on the rRNA (Li *et al*, 2011b, 2011a; Wang & Meier, 2004). While pseudouridylation itself is not essential for yeast, it is required for translational fidelity, which can be easily assessed either by sensitivity towards antibiotics or reporters for translational fidelity (Jack *et al*, 2011). As co-transcriptional pseudouridylation has been observed in human mRNA (Martinez *et al*, 2020), one straight-forward model would have been an Nhp2-mediated recruitment to the rDNA chromatin to place Cbf5 in close proximity to its target. However, we were not able to detect any evidence linking H2AQ105me and pseudouridylation. We cannot rule out the possibility that the assays were not sensitive enough as an H2AQ105A mutant still recruits around 50% of Nhp2 to the rDNA (Figure 4A). However, we did not observe enrichment for members of the Cbf5 catalytic complex on peptide pulldowns arguing that the Nhp2-H2AQ105me interaction is independent of Nhp2’s function in the pseudouridylation complex. Indeed, intersecting binders of H2AQ105me and Nhp2, we identified members of the small subunit processome (SSU) complex to common interactors. Interestingly, also mutating H3K56 to alanine leads to defects in SSU recruitment to the rDNA (Chen *et al*, 2012). As the SSU functions co-transcriptionally (Gallagher *et al*, 2004; Phipps *et al*, 2011), it is tempting to speculate that the Nhp2 recruitment by H2AQ105me and the reported recruitment of the SSU via H3K56ac allow for an efficient coupling of transcription and pre-rRNA processing.

## MATERIALS AND METHODS

### Strains, plasmids and reagents

Genotypes of yeast strains and plasmids are listed in Supplementary Table 4. All chemicals used in this study were purchased analytical grade from either Sigma Aldrich or Carl Roth, unless stated otherwise. Drop Out Mix for yeast synthetic medium was from US Biological Life Sciences (D9543-01). 5-FOA was bought from Cayman Chemicals (17318). Protein-G coupled Dynabeads™ (10004D) and streptavidin-coated Dynabeads™ MyOne™ Streptavidin C1 (65001) were from Thermo Fisher. Secondary antibodies against rabbit (7074S) and mouse IgG (7076S) coupled to HRP were purchased from Cell Signaling. Primary antibodies were as follows: H3K56ac (Active Motif - 39281), G6PDH (Sigma - A9521) and c-Myc (9e10, Sigma - MABE282), FLAG (Sigma - F3165), H2AQ105me from (Tessarz *et al*, 2014), H2A (Abcam - ab13923), H3 (Abcam - ab1791), Utp11L (GeneTex GTX115929), Dyskerin (Santa Cruz sc-373956) and human Nhp2 (Proteintech 15128-1-AP). ECL solution was from Promega (W1001).

### Yeast protocols

If not stated otherwise, all strains used were derived from W303. Gene deletions or tag integrations were performed using PCR-based methods (Janke *et al*, 2004; Longtine *et al*, 1998). Yeast were grown in YPD medium if not stated differently. Histone mutants were generated using plasmid shuffling. For spot tests, cells were grown over-night to stationary phase in YPD, diluted to an optical density (OD_600nm_) of 1 and serially diluted 5-fold before spotting on YPD plates containing the relevant antibiotics. Synthetic genetic array was performed essentially as previously described (Tong *et al*, 2001) using the knock-out (Giaever *et al*, 2002; Breslow *et al*, 2008) and DAMP (Breslow *et al*, 2008) collections.

### Protein precipitation, SDS-PAGE and Western Blot

Protein was extracted from a yeast pellet of OD_600nm_ of 1, using sodium hydroxide lysis and TCA precipitation as previously described (Knop *et al*, 1999). Precipitated proteins were dissolved in 50 µl of 2x Laemmli loading buffer and boiled for 10 minutes. 10 µl were loaded per well of a 15% SDS-PAGE, which was run at 120-200 V in 1x SDS gel running buffer until the bromophenol blue marker reached the bottom of the gel. Separated proteins were either visualised directly using coomassie blue staining or transferred to a nitrocellulose membrane using 1x carbonate buffer (10 mM NaHCO_3_, 3 mM Na_2_CO_3_, pH 9.0, 20% methanol) at 400 mA for 75 min. Membranes were blocked with 5% BSA (for histone and histone modification antibodies) or 5% milk (for all other antibodies in 1x TBST for one hour at room temperature. Primary antibody was added, following manufacturer’s recommended concentration, in blocking solution. Membranes were incubated overnight at 4°C with gentle rocking, subsequently incubated with secondary antibody and developed using ECL.

### Recombinant expression of Nhp2

pET24a-Nhp2 was transformed into BL21 DE3 Codon Plus and plated on LB-Agar, supplemented with kanamycin and chloramphenicol. Overnight cultures were set up and diluted to OD_600nm_ 0.1 in 3x 1.5 liters in 2YT supplemented with kanamycin and chloramphenicol and grown to OD_600nm_ 0.6 at 37°C. 2mM IPTG were added for 3 hours. Cells were harvested and 1 pellet per 1.5 liters was frozen and stored at −80°C. For purification, each pellet was resuspended in 35ml WB 1 (20mM Tris pH 7.5, 300mM NaCl, 5mM β-Mercaptoethanol, 10mM Imidazole) and sonicated on ice using a Branson sonifier 450 with 50% duty cycle and output control 2 for 6 × 30sec with 1min incubation on ice in between. Cell lysates were centrifuged at 20,000g for 30min at 4°C. Lysates were combined, passed through a 0.45μm filter and incubated with 1ml Ni-NTA beads (Qiagen) for 1h at 4°C. Beads were washed with 50ml WB1, followed by 50ml WB2 (20mM Tris pH 7.5, 100mM NaCl, 5mM β-Mercaptoethanol, 10mM Imidazole) and eluted in 1ml fractions using EB1 (20mM Tris pH 7.5, 100mM NaCl, 5mM β-Mercaptoethanol, 250mM Imidazole). Fractions were checked by SDS-PAGE for protein and those containing Nhp2 were pooled and passed over a 2ml SP-sepharose column equilibrated with SP1 (20mM Tris pH 7.5, 100mM NaCl, 5mM β-Mercaptoethanol). Column was washed with 50ml SP1 and Nhp2 was eluted with SP2 (20mM Tris pH 7.5, 1M NaCl, 5mM β-Mercaptoethanol) and fractions of 1ml were collected and checked for Nhp2. Fractions containing pure Nhp2 were pooled and dialyzed against storage buffer (20mM Tris pH 7.5, 150mM NaCl, 5mM β-Mercaptoethanol, 10% glycerol), aliquoted, snap frozen and stored at −80°C.

### Chromatin immunoprecipitation, qPCR or next generation sequencing and analysis

Yeast was grown into mid-log phase, (OD_600nm_ 0.6) in YPD, and 50 ml were harvested and placed into 50ml falcon tubes. 1% formaldehyde (final concentration) was added and cultures were left to rotate at room temperature for 30 minutes to allow for crosslinking. Cultures were centrifuged (3K, 2 min, 4°C) and supernatant was discarded. Cell pellets were resuspended in 20 ml cold PBS, centrifuged as above and the supernatant discarded again. This washing was repeated 2 times in total and cells were then transferred to 2 ml eppendorf tubes and kept on ice. Pellets were then resuspended in 500 µl SDS buffer (1% SDS, 10mM EDTA, 50mM Tris-Cl (pH 8.0), plus protease inhibitors) on ice. About 200 µl of acid-washed glass beads were added and tubes were vortexed at top speed at room temperature for 6x 1 min, with 3 min on ice in between. The bottoms of the tubes were pierced with a hot needle and place in 15ml falcon tubes, which were centrifuged (2k, 30 seconds) to release the cell lysate. Lysates were sonicated to produce fragments of 200-600bp. Sonicated samples were transferred to 1.5 ml Eppendorf tubes, centrifuged (20 min, 14K, 4°C) and diluted in 5 ml of cold IP buffer (0.01% SDS, 1.1%Triton-X-100, 1.2mM EDTA, 16.7MM Tris-Cl (pH 8.0), 167mM NaCl, plus protease inhibitors). 50 µl were taken as input control and stored at −20°C. For the IP, 1 ml of chromatin was placed in siliconized Eppendorf tubes and 2 µg of antibody was added. Chromatin/antibody mix was place on rotating wheel at 4°C overnight. 30 µl of Protein G Dynabeads ™ were washed 3x in IP buffer and added to the chromatin/antibody mix. This was incubated at room temperature on a rotating wheel for 90 minutes. Beads were washed for 3 min with each wash buffer (1 ml TSE (1% Triton-X-100, 0.1%SDS, 2mM EDTA, 20mM Tris-Cl (pH 8.0)) plus 150 mM NaCl, 1ml TSE plus 500 mM NaCl, 1 ml LiCl wash (0.25M LiCl, 1% NP-40, 1% dioxycholate, 1mM EDTA, 10mM Tris-Cl (pH 8.0)), 1 ml TE (pH8.0)). IP was eluted using 200 µl of Elution buffer (1% SDS, 0.1M NaHCO_3_), made fresh on the day, on a rotating wheel for 30min at room temperature. 20 µl of 5M NaCl was added to all samples, which were then incubated at 95°C for 15 min, or overnight at 65°C, to reverse the formaldehyde crosslink. 5 µg of DNase-free RNase was then added for 30 min at 37°C and then the DNA was purified using a Qiagen PCR purification kit. 1 µl was analysed per reaction by qPCR. ChIP-seq libraries were generated following a previously published protocol (Ford *et al*, 2014). A detailed version of this protocol is available upon request.

### ChIP-seq analysis

For ChIP-seq, samples were single-end deep-sequenced and reads were aligned to the sacCer3 genome using Bowtie2 (v 2.0) (Langmead & Salzberg, 2012). Peak calling was performed using Macs2 (Zhang *et al*, 2008) with peaks displaying an FDR < 10^−5^ considered statistically significant. All aligned ChIP-seq BAM files were converted to bigwig (10 bp bin) and normalized to total sequencing depth using deepTools (v 2.0) (Ramírez *et al*, 2016) . Single gene tracks were generated through the IGV genome browser.

### SILAC labelling

BY4742 was grown in 1 L of SD supplemented with amino acids and uracil, excluding lysine. Cultures were then supplemented with 100 mg/L of either normal lysine or heavy lysine (Sigma, L-Lysine-^13^C_6_ ^15^N_2_ hydrochloride cat. number 608041) and grown for no less than 15 hours to reach an OD_600nm_ of 0.6-0.8, to allow for maximum labeling of proteins. Cells were then harvested following the peptide-pulldown protocol.

### Yeast lysates

Cells were grown in 200 ml of YPD (for SILAC labelling see specific culture protocol) and harvest and washed 1x in 1x PBS, to get rid of excess media, and resuspended in a high-salt binding buffer (50mM Tris-Cl (pH 8.0), 1% NP40, 420 mM NaCl, 1mM DTT, and protease inhibitors) 1:2 (yeast:buffer). Yeast suspension was dropped directly into liquid nitrogen to snap freeze and then lysed using a Freezer Mill. Cell debris was removed by centrifugation (10K, 4**°**C, 2 min) and the supernatant was transferred to a clean tube and diluted 1:1 with no-salt binding buffer (50mM Tris-Cl (pH 8.0), 1% NP40, 1mM DTT, and protease inhibitors). Protein content was verified by Bradford assay and the lysate was further diluted, in binding buffer (50mM Tris-Cl (pH 8.0), 1% NP40, 150 mM NaCl, 1mM DTT, and protease inhibitors), to obtain a concentration of about 1.7 mg/ml.

### Pull-downs

600 ul of cleared yeast lysates (about 1 mg of protein) was used per pulldown. For peptide pulldowns, cell lysates were incubated with peptide-coupled magnetic beads Dynabeads ™ MyOne ™ Streptavidin C1. Lysate/bead mix was rotated at 4**°**C for 2 hours and then washed 5x in 1ml of binding buffer. For analysis by western blot, proteins were eluted in 2x Laemmli buffer for 10 minutes at 65**°**C, boiled for a further 10 min, and then loaded onto an SDS PAGE and analysed by western blot. For mass spectrometry see Mass Spectrometry section. Beads were washed 5x with 1ml TBS at 4**°**C before protein elution in mass spec elution buffer. For the Nhp2-myc IP, 5μl anti-myc antibody was bound to 50 μl ProteinG Dynabeads for at least 2h at 4**°**C in TBS. Cleared protein lysates were adjusted to 1mg/ml and 1ml were added to prebound antibodies and incubated at 4**°**C overnight. Beads were washed 5x with 1ml TBS at 4**°**C before protein elution in mass spec elution buffer (see Mass Spectrometry protocol).

### Mass Spectrometry sample preparation

Following peptide pulldown of SILAC-labelled cultures, beads from heavy and light experiments were mixed and washed further 2x in 1 ml of 1x TBS to remove excess detergent. Proteins were eluted in 50ul of mass spec elution buffer (6M Guanidine hydrochloride, 10mM TCEP, 40mM CAA, 100mM Tris (pH 8.5)), for 1 hour at room temperature, regularly shaking to prevent the beads from settling, and then diluted 10-fold in 20 mM Tris/10 % acetronile. The protein concentration was measured using a nanodrop (260/280) and 5 ug of LysC was added to 100 ug of protein. Proteins were digested on the beads at 37**°**C overnight with gentle shaking so as to prevent the beads from settling. Following protein digestion, peptides were purified using FASP peptide purification (Coleman *et al*, 2017), followed by Mass spec analysis.

### LC-MS/MS analysis

Peptides were separated on a 25 cm, 75 μm internal diameter PicoFrit analytical column (New Objective) packed with 1.9 μm ReproSil-Pur 120 C18-AQ media (Dr. Maisch,) using an EASY-nLC 1000 (Thermo Fisher Scientific). The column was maintained at 50°C. Buffer A and B were 0.1% formic acid in water and 0.1% formic acid in acetonitrile. For the Qme IP, peptides were separated on a segmented gradient from 5% to 20% buffer B for 100 min, from 20% to 25% buffer B for 10 min, and from 25% to 40% buffer B for 10 min at 200 nl / min. For the Nhp2 IP peptides were separated on a segmented gradient from 6% to 31% buffer B for 45 min, and from 31% to 44% buffer B for 8 min at 200 nl / min. Eluting peptides were analyzed on a QExactive Plus (Qme IP) or QExactive HF (Nhp2 IP) mass spectrometer (Thermo Fisher Scientific). Peptide precursor m/z measurements were carried out at 70000 (Qme IP) or 60000 (Nhp2 IP) resolution in the 300 to 1800 (Qme IP) or 300 to 1500 (Nhp2 IP) m/z range. The top ten most intense precursors with charge state from 2 to 7 only were selected for HCD fragmentation using 25% (Qme IP) or 27% (Nhp2 IP) normalized collision energy. The m/z values of the peptide fragments were measured at a resolution of 17500 (Qme IP) or 15000 (Nhp2 IP) using 80 ms maximum injection time. Upon fragmentation, precursors were put on a dynamic exclusion list for 45 sec.

### Protein Identification and Quantification

The raw data were analyzed with MaxQuant version 1.5.2.8 (Cox & Mann, 2008) using the integrated Andromeda search engine (Cox *et al*, 2011). Peptide fragmentation spectra were searched against the canonical and isoform sequences of the yeast reference proteome (proteome ID UP000002311, downloaded February 2015 from UniProt). Methionine oxidation and protein N-terminal acetylation were set as variable modifications; cysteine carbamidomethylation was set as fixed modification. The digestion parameters were set to “Specific” and “LysC/P” (Qme IP) or “Trypsin/P” (Nhp2 IP). The minimum number of peptides and razor peptides for protein identification was 1; the minimum number of unique peptides was 0. Protein identification was performed at a peptide spectrum matches and protein false discovery rate of 0.01. The “second peptide” option was on. For the analysis of the Qme IP, “Re-quantify” was enabled. . For the analysis of the Nhp2 IP, successful identifications were transferred between the different raw files using the “Match between runs” option. SILAC quantification (Qme IP) was performed using a minimum ratio count of two. Label-free quantification (Nhp2 IP) (Cox *et al*, 2014) was performed using a minimum ratio count of two. Downstream data transformation, filtering and differential abundance analysis was performed using Perseus version 1.5.0.0 (Tyanova *et al*, 2016). For the Qme IP data, log2 transformed SILAC ratios were analyzed using a one-sided t-test against zero. For the Nhp2 IP data, label-free intensities were filtered for at least three valid values in at least one group and imputed from a normal distribution with a width of 0.3 and down shift of 1.8. Imputed values were analyzed with a two-sided t-test, using a S0 parameter of one.

## DATA AVAILABILITY

The mass spectrometry proteomics data have been deposited to the ProteomeXchange Consortium via the PRIDE (Perez-Riverol *et al*, 2019) partner repository with the dataset identifier XXX. Analyzed data can be found in the supplementary material. ChIP-seq data of Nhp2 available at XXX

## ACKNOWLEDGEMENTS

We would like to thank all members of the Tessarz laboratory for discussion and C. Nikopoulou for critical reading of the manuscript. Part of this work was initiated in the laboratory of Tony Kouzarides. We thank Martin Graef for help with the SGA. Mass spectrometry was performed in the Proteomics facility of the Max Planck Institute for Biology of Ageing. Xinping Li advised on sample preparation and acquired mass spectrometry proteomics data, Ilian Atanassov performed proteomics data analysis. ChIP-sequencing was done at the Genomic Core Facility of the MPI for Plant Breeding Research, Cologne, Germany. This work was funded by the Max Planck Society and the German Research Council (to P.T., TE1079/2-1).

## SUPPLEMENTARY FIGURES

**Supplementary Figure 1:**
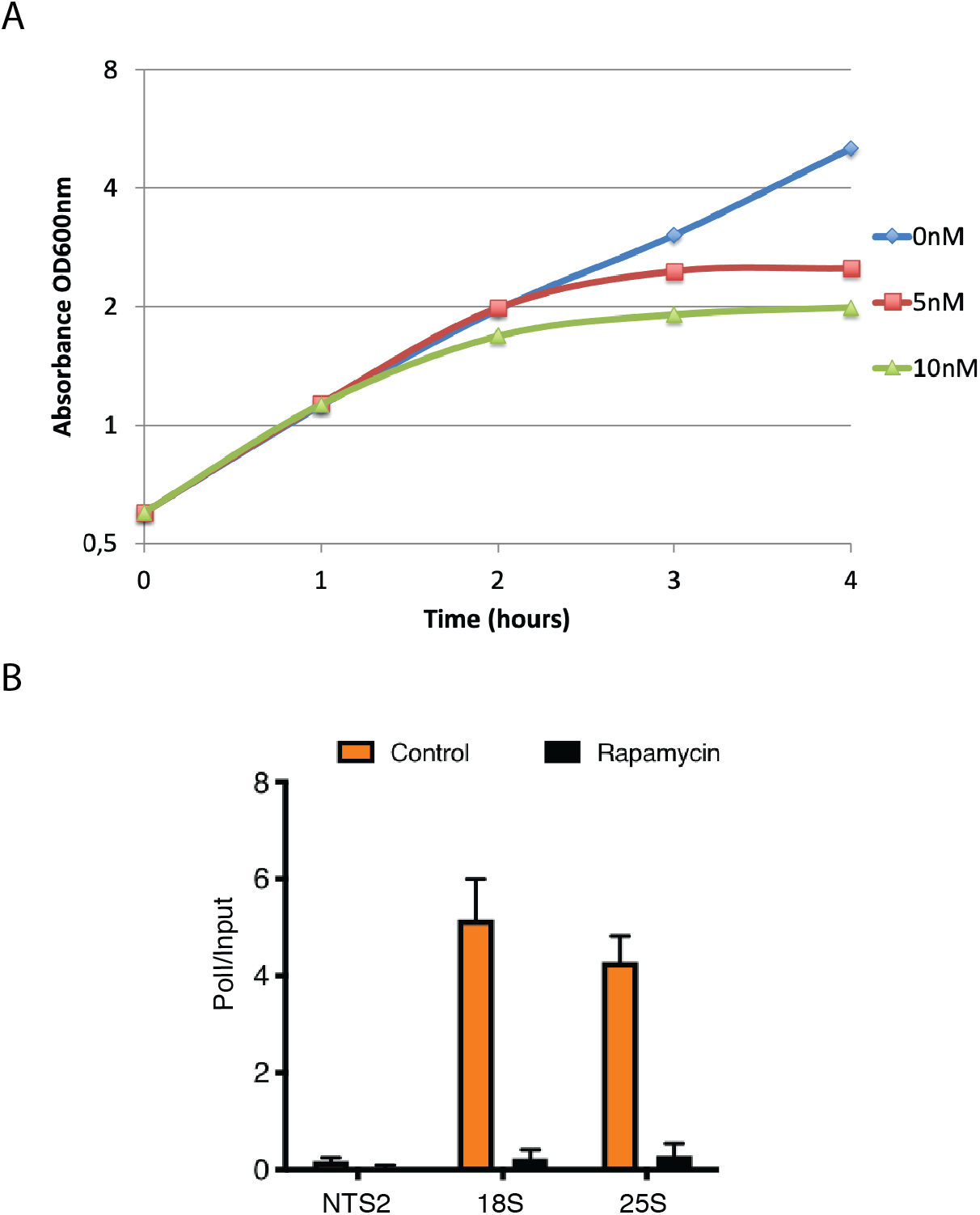
A) Growth curve of WT yeast cells challenged with indicated concentrations of rapamycin. B) Loss of RNA Pol I occupancy at the rDNA upon treatment with 5nM rapamycin.

**Supplementary Figure 2:**
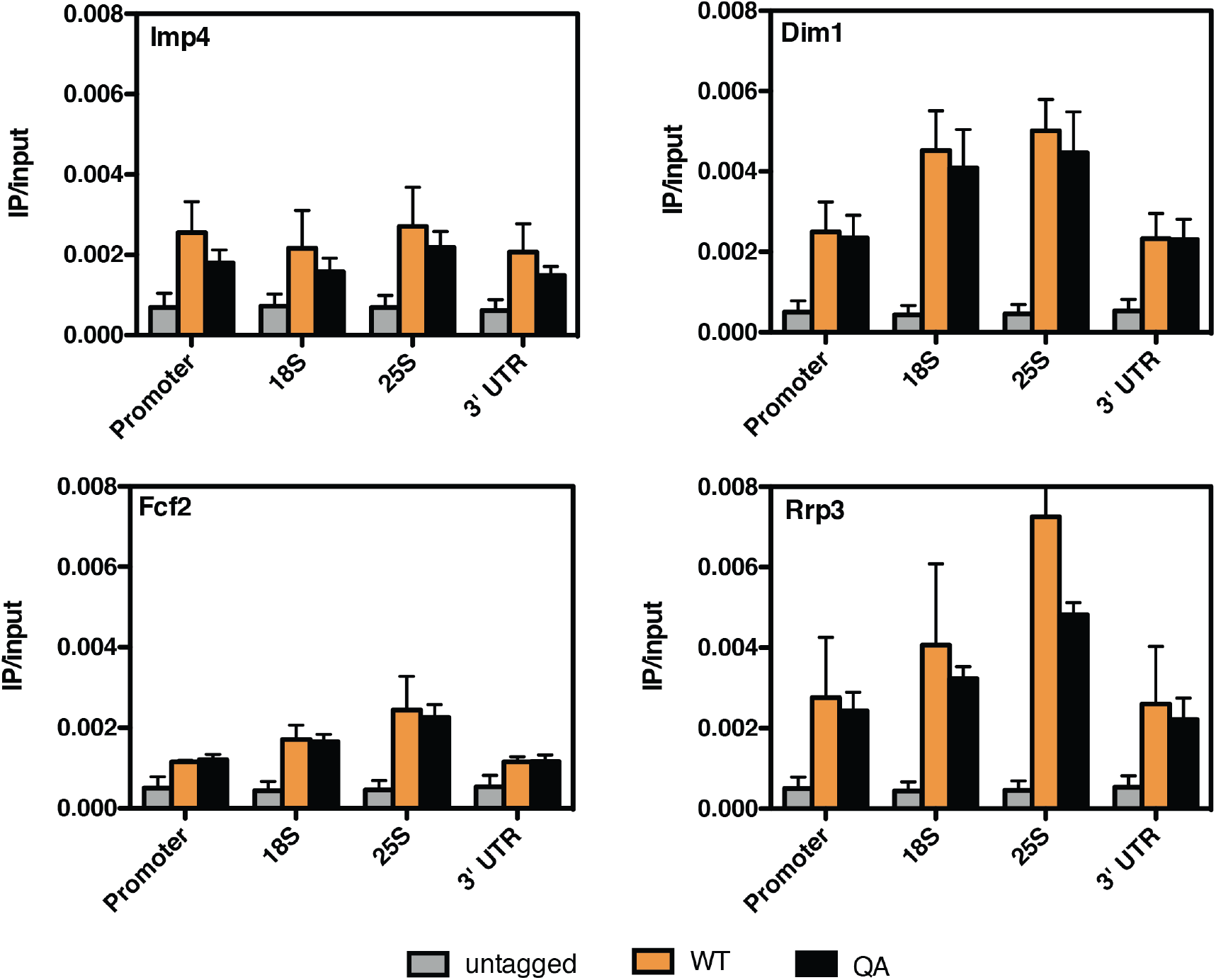
ChIP-qPCR analysis of potential and indicated readers of H2AQ105me based on peptide pulldown.

## Notes

### Competing Interest Statement

The authors have declared no competing interest.

